# Rapid single-step affinity purification of HA-tagged mitochondria from *Arabidopsis thaliana*

**DOI:** 10.1101/672972

**Authors:** Franziska Kuhnert, Anja Stefanski, Nina Overbeck, Kai Stühler, Andreas P.M. Weber

**Author notes:** **Author contributions**: F.K. generated transgenic lines harboring affinity-tagged mitochondria, isolated mitochondria, analyzed the data, and drafted the manuscript; A.S., N.O., and K.S. performed the proteomics analysis and analyzed the data; A.P.M.W. conceived and supervised the experiments and contributed to the writing of the manuscript. **Funding**: This work was supported by the Deutsche Forschungsgemeinschaft (CRC 1208 and funding under Germany’s Excellence Strategy – EXC-2048/1 – project ID 390686111”).

## Abstract

Photosynthesis in plant cells would not be possible without the supportive role of mitochondria. However, isolation of mitochondria from plant cells, for physiological and biochemical analyses, is a lengthy and tedious process. Established isolation protocols require multiple centrifugation steps and substantial amounts of starting material. To overcome these limitations, we tagged mitochondria in *Arabidopsis thaliana* with a triple haemagglutinin-tag for rapid purification via a single affinity purification step. This protocol yields a substantial quantity of highly pure mitochondria from 1 g of Arabidopsis seedlings. The purified mitochondria were suitable for enzyme activity analyses and yielded sufficient amounts of proteins for deep proteomic profiling. We applied this method for the proteomic analysis of the Arabidopsis *bou-2* mutant deficient in the mitochondrial glutamate transporter À bout de souffle (BOU) and identified 27 differentially expressed mitochondrial proteins compared with transgenic Col-0 controls. Our work also sets the stage for the development of advanced mitochondria isolation protocols for distinct cell types.

**One-sentence summary:** Affinity-tagging of mitochondria in plant cells with a triple hemagglutinin-tag enables single-step affinity purification of mitochondria in less than 20 min.

## INTRODUCTION

In all eukaryotic organisms, mitochondria are the major source of ATP, which is produced via the oxidative phosphorylation (OXPHOS) pathway, thus playing a vital role in cellular energy metabolism. Mitochondria also participate in amino acid metabolism as well as in photorespiration in photosynthetic eukaryotes. Photorespiration plays a crucial role in photosynthesis by detoxifying 2-phosphoglycolate, which is produced by the oxygenation of Rubisco and acts as an inhibitor of several plastidial enzymes (Ogren and Bowes, 1971; Kelly and Latzko, 1976; Husic et al., 1987). Plants reclaim 2-phosphoglycolate in the complex pathway of photorespiration, yielding 3-phosphoglycerate, which is returned to the Calvin Benson cycle. The photorespiratory pathway includes several enzymatic steps that occur in four subcellular compartments: plastids, peroxisomes, mitochondria, and cytosol (Eisenhut et al., 2019). Knockout mutants of genes encoding enzymes and transporters involved in photorespiration often show a photorespiratory phenotype, characterized by chlorotic leaves and growth inhibition under ambient carbon dioxide (CO_2_) conditions, which can be rescued in a CO_2_ enriched environment (Peterhansel et al., 2010). A key step in photorespiration is the conversion of two glycine molecules into one serine residue in the mitochondrial matrix, accompanied by the release of CO_2_ and ammonia. This step is catalyzed by the glycine decarboxylase (GDC) multienzyme system, comprising a P-protein (GLDP), H-protein (GDCH), L-protein (GDCL), and T-protein (GLDT), in combination with serine hydroxymethyltransferase (SHM) (Voll et al., 2006; Engel et al., 2007). In green tissues, these proteins constitute up to 50% of the total protein content of the mitochondrial matrix, indicating the importance of glycine oxidation in mitochondria (Oliver et al., 1990).

The *Arabidopsis thaliana bou-2* mutant was previously identified as lacking the mitochondrial glutamate transporter À bout de souffle (BOU), which is involved in photorespiration (Eisenhut et al., 2013). Plants lacking the inner mitochondrial membrane (IMM) protein BOU show a pronounced photorespiratory phenotype under ambient CO_2_ conditions, significantly elevated CO_2_ compensation point, and highly reduced GDC activity in the isolated mitochondria (Eisenhut et al., 2013). Because *BOU* is co-expressed with genes encoding components of the GDC complex, and the *bou-2* mutant shows a similar metabolic phenotype as the *shm1* mutant, it was hypothesized that BOU transports a metabolite necessary for the proper functioning of GDC (Voll et al., 2006; Eisenhut et al., 2013). Recently, it was demonstrated that heterologously expressed BOU functions as a glutamate transporter (Porcelli et al., 2018). Glutamate is neither a substrate nor a product of the reaction catalyzed by GDC. Besides its role in amino acid and N metabolism, glutamate is necessary for the glutamylation of tetrahydrofolate (THF), a cofactor of GLDT and SHM (Suh et al., 2001). Glutamylation of THF increases its stability. Moreover, THF-dependent enzymes generally prefer polyglutamylated folates over monoglutamylated folates as a substrate (Suh et al., 2001). However, because (1) BOU is not the only glutamate transporter in mitochondria, and (2) glutamylation of folates is not restricted to mitochondria, the exact physiological function of BOU remains unclear (Hanson and Gregory, 2011; Monné et al., 2018). Notably, a glutamate/glutamine shuttle across the mitochondrial membrane was previously suggested to support the reclamation of ammonia released during photorespiration (Linka and Weber, 2005).

Analysis of the biochemical and physiological functions of mitochondria frequently requires the isolation of intact mitochondria. Mitochondria can be isolated from leaf tissue in less than 1 h by differential centrifugation. This method yields mitochondria with good integrity and appropriate enzyme activity. However, mitochondria are frequently contaminated with plastids and peroxisomes. Hence, in many cases, a combination of differential centrifugation and Percoll density gradient is used (Millar et al., 2001; Werhahn et al., 2001; Keech et al., 2005). While this produces a pure fraction of respiratory active mitochondria with low plastid and peroxisome contamination, such procedures generally take several hours and require up to 50 g of starting material for producing sufficient yields (Keech et al., 2005). Moreover, traditional protocols are not practical for the isolation of mitochondria from mutants with severely impaired growth or from less abundant tissue types such as flowers and for the analysis of mitochondrial metabolites. Furthermore, media used for the isolation of mitochondria typically contain high concentrations of sugars and other metabolites that can potentially interfere with MS-based metabolite analyses.

Recently, Chen and coworkers reported a method for the rapid isolation of mitochondria from human HeLa cell cultures via co-immunopurification (co-IP) (Chen et al., 2016). The authors generated transgenic HeLa cell lines expressing a triple hemagglutinin (HA)-tagged enhanced green fluorescent protein (eGFP) fused to the outer mitochondrial membrane (OMM) localization sequence of OMP25 (3×HA-eGFP-OMP25). Because the epitope-tag was displayed on the surface of mitochondria, these transfected cell lines could be used to rapidly enrich mitochondria after cell homogenization. The HA-tagged mitochondria were captured and pulled down using magnetic beads coated with an anti-HA-tag antibody. Given the small size (1 µm diameter) and non-porous behavior of anti-HA-tag beads, these beads performed better than the porous agarose matrix for the enrichment of mitochondria. Thus, the authors established a method that ensures a high yield of pure mitochondria in approximately 12 min. The isolated mitochondria showed high purity, integrity, and functionality. Additionally, the authors developed a simple potassium-based buffer system that maintains mitochondrial intactness and is compatible with downstream analyses, such as metabolite analysis by LC/MS (Chen et al., 2016).

Building on this previous work, we developed an affinity-tagging strategy for the rapid isolation of mitochondria from Arabidopsis. We generated transgenic Arabidopsis lines carrying an HA-tagged translocase of the OMM 5 (TOM5) and isolated highly pure and intact mitochondria from these lines in less than 25 min. The isolated mitochondria were successfully subjected to proteomics and enzyme activity analyses. Moreover, we applied the isolation strategy to the *bou-2* mutant, revealing differential protein abundance and enzyme activities.

## RESULTS

### Identification of TOM5 a suitable anchor peptide and generation of affinity-tagged Arabidopsis lines

Recently, Chen and colleagues reported a rapid protocol for the isolation of intact mitochondria from transgenic HeLa cells expressing the mitochondrial fusion protein 3×HA-eGFP-OMP25 using co-IP. The Arabidopsis genome does not encode an ortholog of OMP25. Therefore, we screened the available Arabidopsis mitochondrial proteome data, including the OMM proteins with known topology and function, and identified TOM5 as a potential candidate for our mitochondria affinity-tagging approach. Together with TOM6, TOM7, TOM20, TOM22/9, and TOM40, TOM5 forms the protein import apparatus of plant mitochondria (Werhahn et al., 2003). In yeast (*Saccharomyces cerevisiae*), TOM5 is an integral protein of the OMM. It has a negatively-charged N-terminal domain, which faces toward the cytosol (Dietmeier et al., 1997) and can be fused to GFP without altering the subcellular localization of TOM5 (Horie et al., 2003). The protein import machinery is well conserved among eukaryotes. The predicted N-terminal cytosolic domain of Arabidopsis TOM5 is necessary for the recognition of cytosolically synthesized mitochondrial preproteins (Wiedemann et al., 2004). Therefore, we generated an N-terminal translational fusion of the Arabidopsis *TOM5* gene with triple HA-tagged *synthetic GFP* (*sGFP*) gene under the control of the Arabidopsis *UBIQUITIN10* promoter (*UB10p*) (*UB10p-3×HA-sGFP-TOM5*; Supplemental Fig. S1). The construct was used to stably transform Arabidopsis ecotype Columbia (Col-0) and *bou-2* mutant. Expression and localization of the fusion protein was verified in root and leaf tissues of 10-day-old Arabidopsis seedlings via confocal laser scanning microscopy. Transgenic Arabidopsis lines expressing the 3×HA-sGFP-TOM5 protein in leaf and root mitochondria of Col-0 and *bou-2* seedlings were identified based on the co-localization of the fluorescent signal of GFP signal with that of the mitochondrial marker CMXRos (Fig. 1A–D). CMXRos is a lipophilic cationic dye that accumulates in the mitochondria because of the negative membrane potential; thus, it solely stains mitochondria with an intact respiratory chain (Pendergrass et al., 2004). Because the fluorescent signals of the 3×HA-sGFP-TOM5 protein (green) fully overlapped with those of the MitoTracker (red), we conclude that overexpression of the *UB10p-3×HA-sGFP-TOM5* construct in Col-0 and *bou-2* does not affect mitochondrial intactness. In a few transgenic Col-0 lines, the fluorescent signal of sGFP formed a ring around the mitochondria, suggesting the localization of 3×HA-sGFP-TOM5 to the OMM (Fig. 1A, B). Notably, transgenic Col-0 or *bou-2* lines showed no apparent phenotypic differences compared with non-transgenic Col-0 or *bou-2* plants (control), respectively, under our culture conditions.

**Figure 1:**
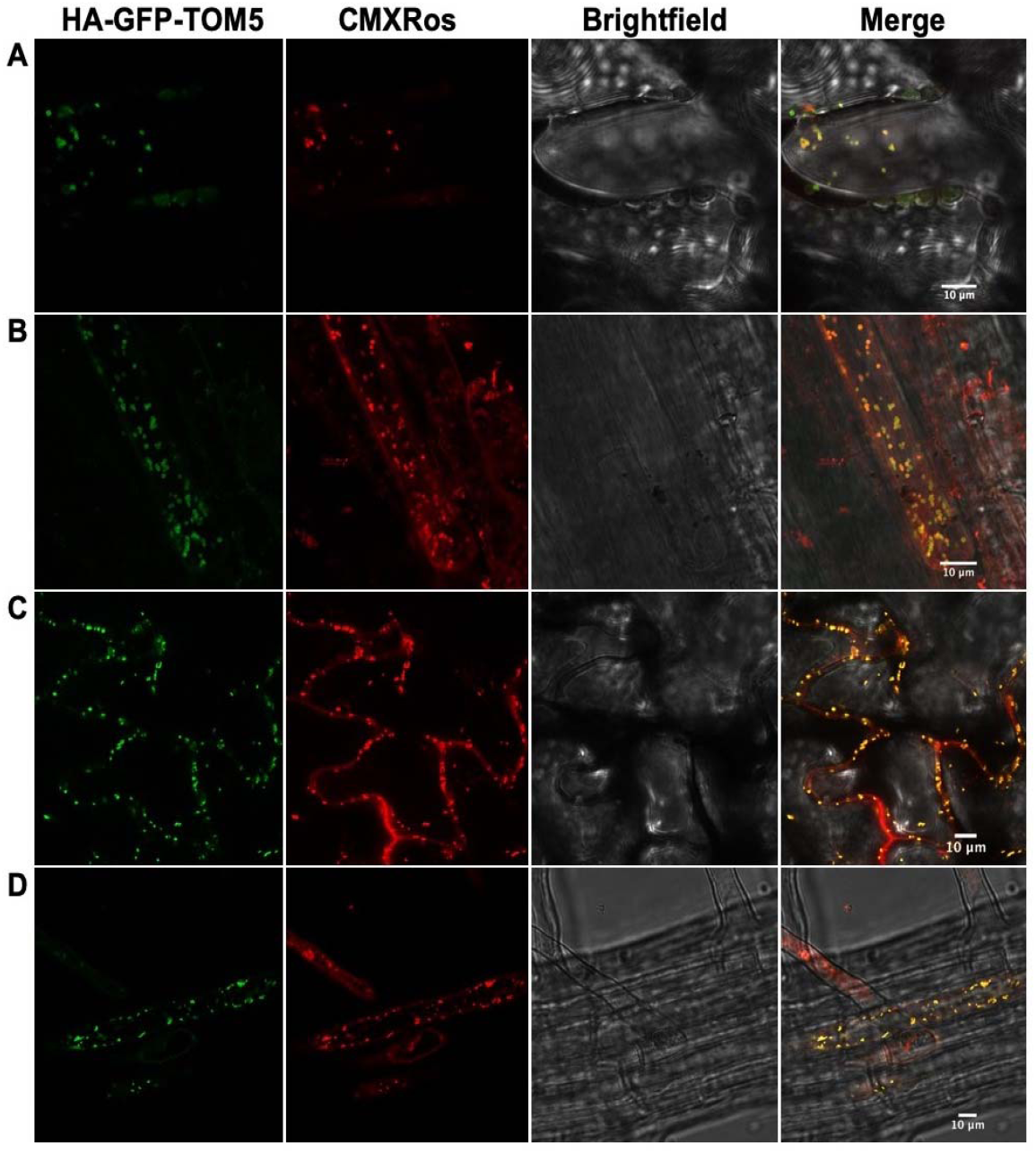
Confocal microscopy of epitope-tagged mitochondria in leaf and root tissues of transgenic Arabidopsis Col-0 and *bou-2* lines expressing the *UB10p-3×HA-sGFP-TOM5* construct. (A–D) Images of transgenic Col-0 leaf (A) and root (B) tissues and transgenic *bou-2* leaf (C) and root (D) tissues expressing the 3×HA-sGFP-TOM5 protein. Green color represents GFP signal, whereas red color represents the signal of mitochondrial marker MitoTracker™ Red CMXRos. Bright field and merged images are shown in yellow. Scale bar = 10 µm.

### Affinity-based purification of intact mitochondria

Transgenic Col-0 and *bou-2* lines were used for the isolation of intact HA-tagged mitochondria via co-IP using magnetic anti-HA beads. We chose HA as the epitope-tag for purification because it has a high affinity for its cognate antibody, and Chen and coworkers previously demonstrated that the size and non-porous behavior of the anti-HA beads yields a high amount of mitochondria (Chen et al., 2016). Additionally, we used the LC/MS-compatible buffer containing KCl and KH_2_PO_4_ (KPBS) developed by Chen et al. (2016). Our purification procedure included five steps: homogenization of plant material in KPBS (1 min), filtration of the homogenate (1 min), two centrifugation steps (5 min and 9 min), and co-IP (7 min, including washing steps). Altogether, mitochondria were purified from plant material in less than 25 min (Fig. 2). If the first centrifugation step used to remove contaminating chloroplasts and cell debris is omitted, the isolation time can be reduced to 18 min. The purified mitochondria were verified by immunoblot analyses using known organelle-specific protein markers. Mitochondria were enriched via co-IP only from lines harboring the mitochondrial 3×HA-sGFP-TOM5 protein, as demonstrated by immunoblot analyses with antibodies directed against different mitochondrial marker proteins including isocitrate dehydrogenase (IDH; mitochondrial matrix), alternative oxidase 1/2 (AOX1/2; IMM), and voltage-dependent anion channel 1 (VDAC1; OMM). No enrichment of mitochondria was observed in control Col-0 lines, indicating that the beads bind specifically to the HA-tag on mitochondria in transgenic lines (Fig. 3). Comparison with classical mitochondria isolation protocols using differential centrifugation and density gradient purification revealed that the mitochondrial fraction enriched using our affinity-tagging method showed significantly less contamination with proteins from plastids, peroxisomes, endoplasmatic reticulum (ER), nuclei, and cytosol (Fig. 3). The following proteins were used as markers for different organelles: Rubisco large subunit (RbcL; plastid), catalase (Cat; peroxisome), lumenal-binding protein 2 (BiP2; ER), histone H3 (nucleus), and heat shock cognate protein 70 (HSC70; cytosol).

**Figure 2:**
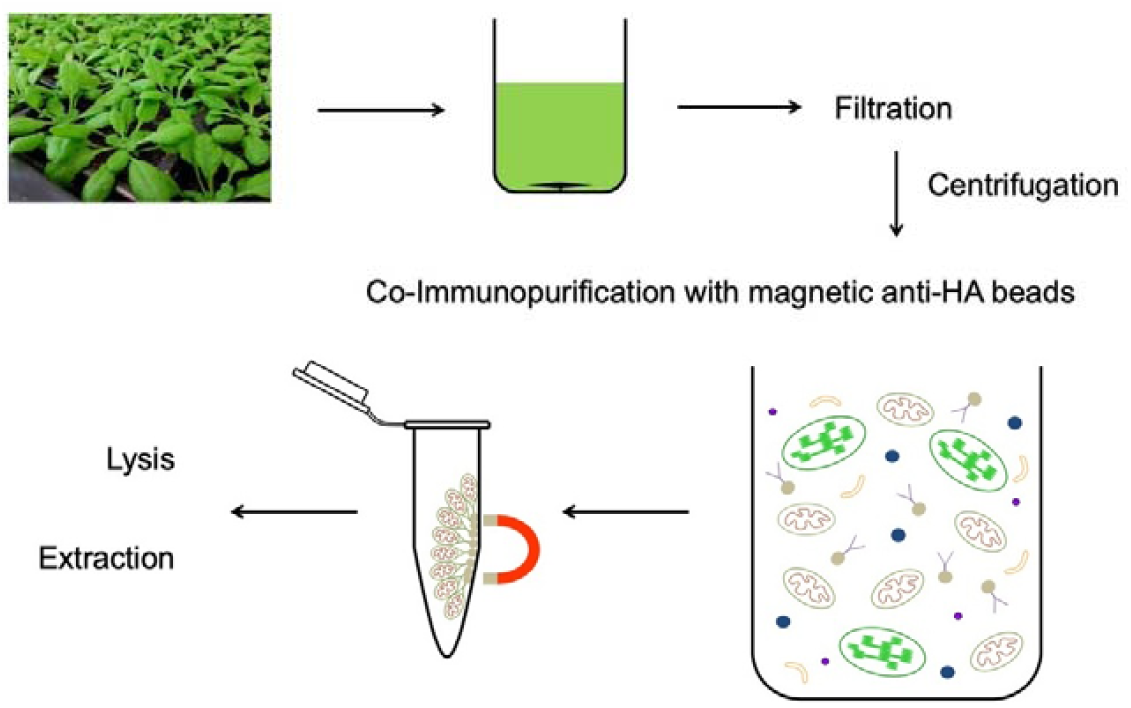
Workflow showing the rapid isolation of epitope-tagged mitochondria via co-Immunopurification (co-IP) from Arabidopsis. Transgenic lines harboring the *UB10p-3×HA-sGFP-TOM5* construct and non-transgenic (control) plants were harvested and homogenized in a Warren blender. The extract was filtered and centrifuged to obtain a crude mitochondrial fraction. Epitope-tagged mitochondria were purified via co-IP using magnetic anti-HA beads. The purified mitochondria were washed and either lysed for immunoblot analysis or extracted for proteomics.

**Figure 3:**
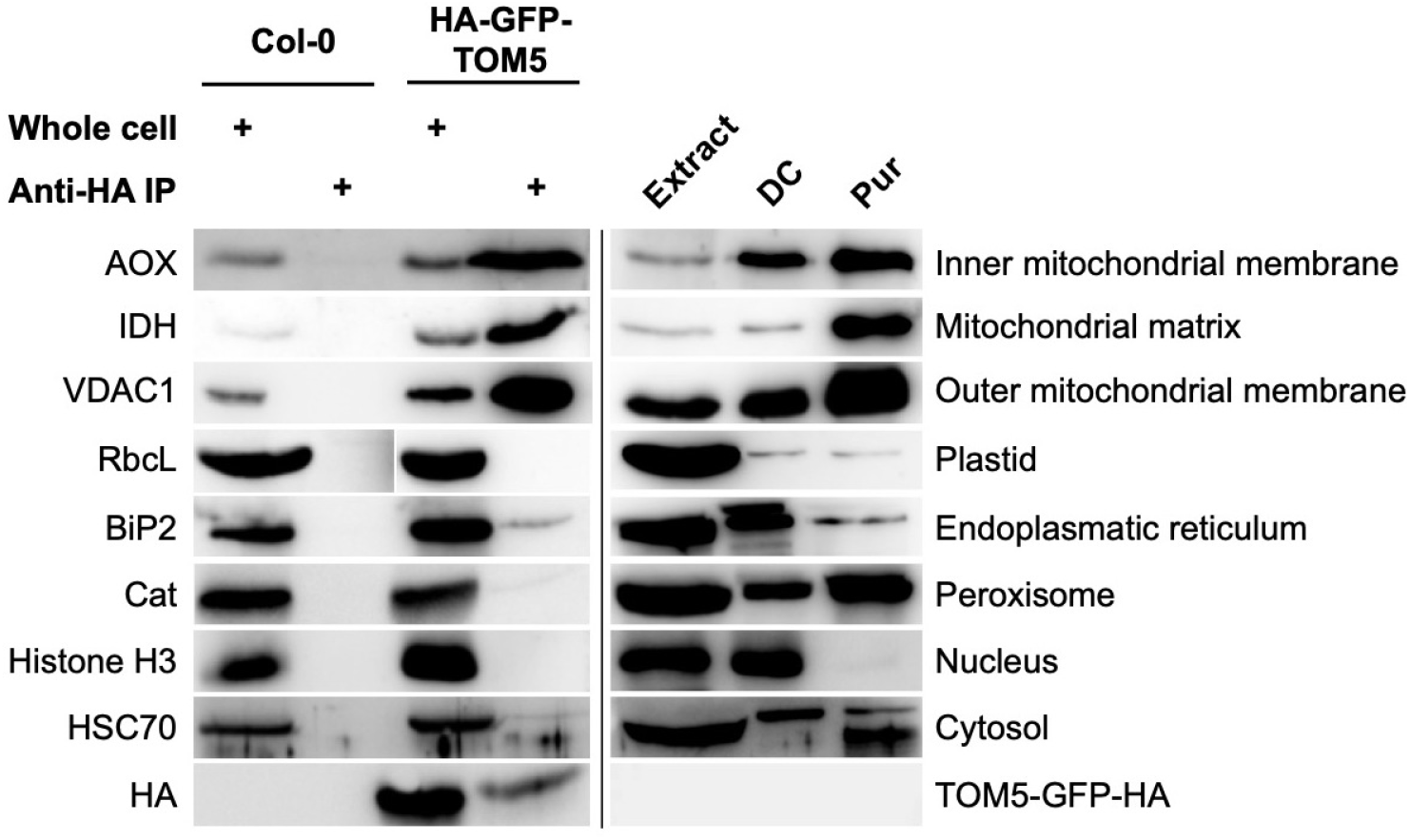
Immunoblot analysis of mitochondria isolated via co-IP or by differential centrifugation and gradient purification. Mitochondria were isolated from transgenic and non-transgenic (control) Col-0 and *bou-2* lines via co-IP using magnetic anti-HA beads (whole cell, Anti-HA IP) or using differential centrifugation and gradient purification (Extract, DC, Pur). Protein amounts were loaded as described in Material and Methods. Whole cell and extract samples were collected after tissue homogenization. Names of organelle marker proteins are shown on the left side of the blots, and their subcellular localization is indicated on the right.

The intactness of mitochondria isolated from HA-tagged and control lines via co-IP was assessed based on the latency of malate dehydrogenase (MDH) activity. Additionally, isolation was performed in the presence of 1% (w/v) Triton X-114. MDH activity could be detected in mitochondria isolated from HA-tagged lines but not in those isolated from the control lines, thus verifying the results of our immunoblot analyses. When 1% (w/v) Triton X-114 was added to the washing buffer during co-IP, MDH activity was undetectable in both transgenic and control lines, as the detergent lyses the organelles (Fig. 4).

**Figure 4:**
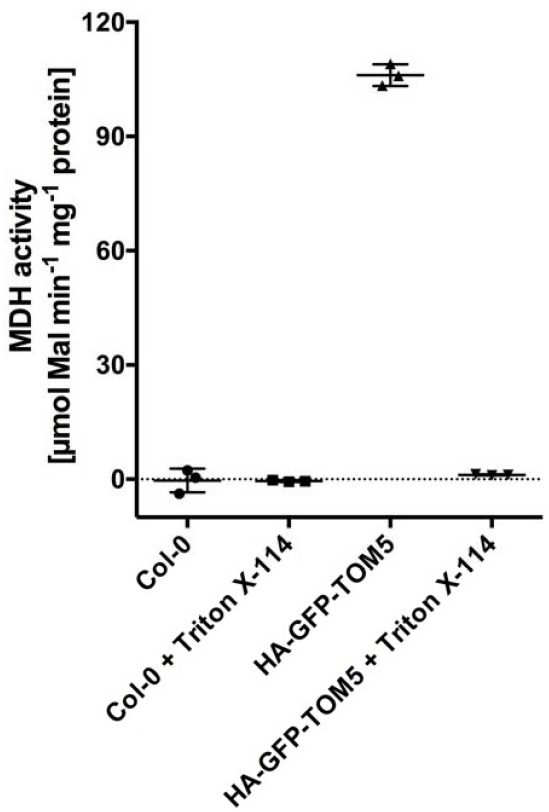
Latency of MDH activity in affinity-purified mitochondria. Mitochondria were rapidly isolated from transgenic and control Col-0 and *bou-2* plants via co-IP, with or without the addition of Triton X-114. The activity of MDH was calculated from the initial slope. Data represent mean ± SD of three replicates.

Typically, we used 5–10 g of Arabidopsis seedlings grown on agar plates for the isolation of mitochondria, and this yielded up to 700 µg of total mitochondrial protein. However, mitochondria could also be isolated from 1 g of starting material, yielding up to 200 µg of total mitochondrial protein. This is advantageous for very young seedlings or mutants with severely impaired growth. Isolation from less than 1 g of starting material may also result in an appropriate yield of mitochondria, but this was not tested in the current study.

Taken together, our data indicate that mitochondria can be rapidly isolated via co-IP using a simple LC/MS-compatible buffer.

### Enzyme assays using mitochondria affinity-purified from transgenic Col-0 and *bou-2* mutant lines

We used our rapid mitochondria isolation method to study the effect of the mitochondrial carrier protein BOU on mitochondrial metabolism in Arabidopsis.

To assess the effect of the *bou-2* mutant allele on mitochondrial metabolism, we rapidly isolated mitochondria from 10-day-old transgenic Col-0 and *bou-2* seedlings grown under elevated CO_2_ conditions (3,000 ppm) and those shifted to ambient CO_2_ conditions (380 ppm CO_2_ after 5 days). Mitochondria were lysed and used to measure the activity of MDH, aspartate aminotransferase (AspAT), glutamate dehydrogenase (GluDH), alanine aminotransferase (AlaAT), γ-aminobutyric acid transaminase (GABA-T), and formate dehydrogenase (FDH). Enzyme activities were calculated from the initial slopes. Enzyme activities in transgenic *bou-2* mutant lines were compared with those in transgenic Col-0 lines, which were set to 100%.

The activities of MDH and FDH were not affected in transgenic *bou-2* mutant lines under any of the conditions tested (Fig. 5A, B). The activities of AspAT and GABA-T in transgenic mutant lines were similar to those in transgenic Col-0 lines, when mitochondria were isolated from seedlings grown under elevated CO_2_ conditions, but were significantly reduced in transgenic mutant lines after the shift to ambient CO_2_ conditions (Fig. 5C, D). The activity of AlaAT was significantly reduced (Fig. 5E), whereas that of GluDH was significantly increased in transgenic mutant lines under elevated CO_2_ conditions; however, both enzyme activities reverted back to the levels in transgenic Col-0 when seedlings were shifted to ambient CO_2_ conditions (Fig. 5F). Together, our results suggest a possible involvement of BOU in mitochondrial amino acid and N metabolism.

**Figure 5:**
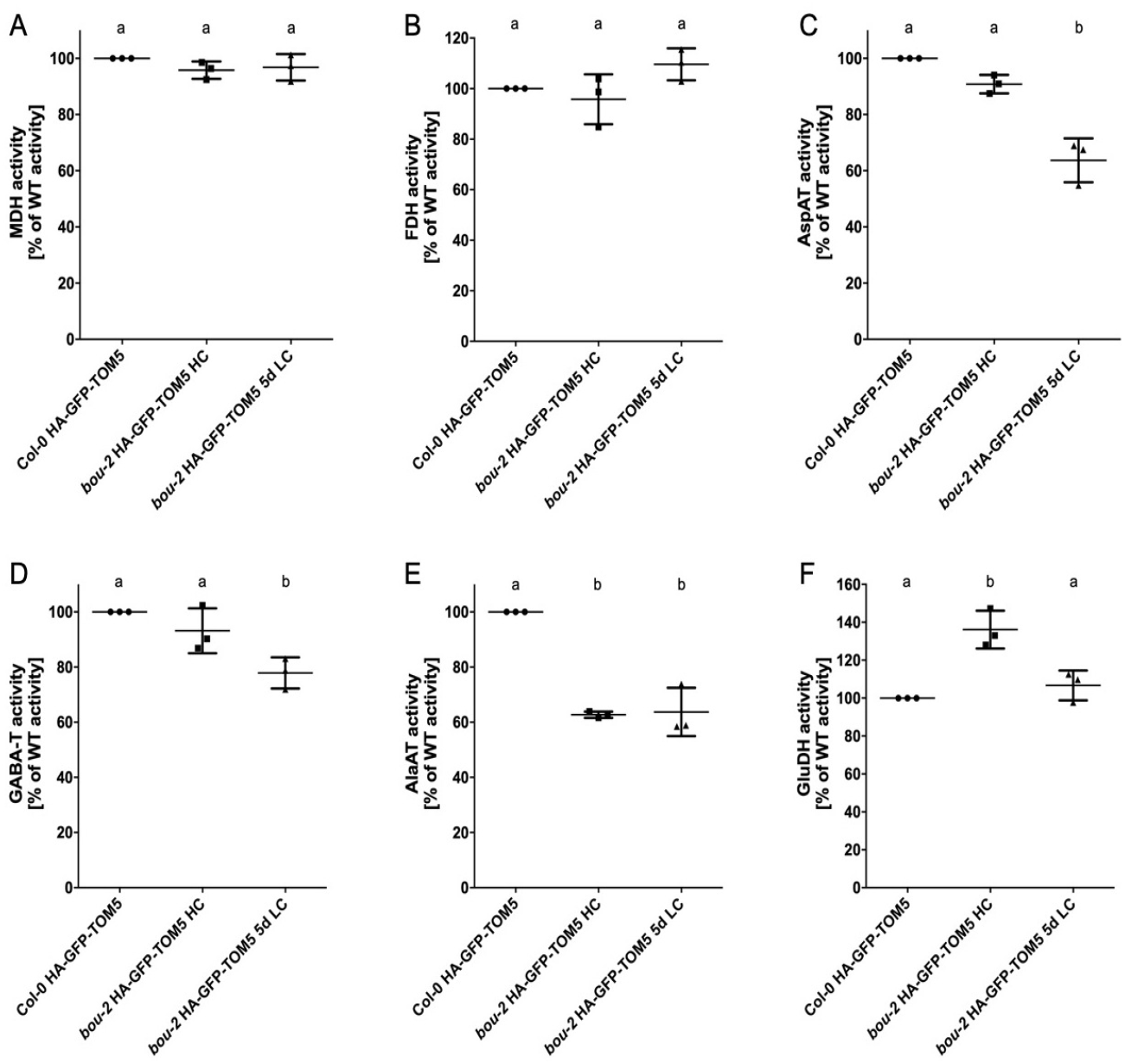
Characterization of enzyme activities in mitochondria isolated from 10-day-old transgenic Col-0 and *bou-2* lines grown at 3,000 ppm (HC) and after shift to 380 ppm (5d LC) via co-IP. (A) Malate dehydrogenase (MDH) activity. (B) Formate dehydrogenase (FDH) activity. (C) Aspartate aminotransferase (AspAT) activity. (D) γ-aminobutyric acid transaminase (GABA-T) activity. (E) Alanine aminotransferase (AlaAT) activity. (F) Glutamate dehydrogenase (GluDH) activity. Activities were calculated from initial slopes. Enzyme activities in transgenic Col-0 were set to 100%. Data represent mean ± SD of three biological replicates. Different letters indicate statistically significant differences between means for each enzyme (*P* < 0.05; one-way ANOVA).

In addition, we found that the activity of MDH was strongly reduced in 4-week-old transgenic *bou-2* mutant plants grown under elevated CO_2_ conditions (Supplemental Fig. S2). However, no change was observed in MDH activity in 10-day-old transgenic *bou-2* plants, suggesting pleiotropic effects in older leaf tissues due to accumulating photorespiratory intermediates.

### Proteomic analysis of mitochondria affinity-purified from transgenic Col-0 and *bou-2* mutant lines

Mitochondria were isolated from 10-day-old transgenic Col-0 and *bou-2* seedlings grown under elevated CO_2_ conditions in four independent biological replicates. Proteome analysis of the isolated mitochondria revealed 15,688 peptides belonging to 1,240 proteins present in at least three of the four replicates (Supplemental Table S1, Supplemental Table S2).

Subcellular localization of the quantified proteins was annotated using the SUBAcon database (Hooper et al., 2017). Summing up the label-free quantitation (LFQ) intensities of the spectra showed that 80% of all quantified peptides resulted from proteins localized or predicted to be localized to the mitochondria, 11.5% from plastid-localized proteins, and 5.7% from proteins with no clear subcellular localization (designated as ambiguous). Additionally, more than 90% of the quantified peptides were assigned to the mitochondria. Contamination of the mitochondrial proteins by proteins from peroxisomes, ER, Golgi, vacuoles, endomembranes, plasma membranes, nuclei, and cytosol was less than 1% each (Supplemental Table S1, Supplemental Table S2). These results indicate high purity of the rapidly isolated mitochondria, which was comparable with the purity of classically isolated mitochondria (Klodmann et al., 2011; Senkler et al., 2017).

Next, we compared our proteome data with previous proteomic analyses and quantified proteins and some of the subunits of known mitochondrial complexes (Supplemental Table S1, Supplemental Table S2). The OXPHOS pathway of mitochondria consists of five protein complexes (I–V) located in the IMM. Complexes I to IV represent oxidoreductases, which comprise the respiratory chain that regenerates oxidized forms of cofactors involved in mitochondrial metabolism, thereby creating an electron flow. This leads to the simultaneous export of protons into the intermembrane space (IMS). The built-up proton gradient is used by Complex V to generate ATP. Complex I is the largest complex involved in the OXPHOS pathway and comprises at least 47 protein subunits that form the so-called membrane and peripheral arms (Peters et al., 2013; Meyer et al., 2019). Except NDUA1, NDUB2, Nad4L, and Nad6 (At3g08610, At1g76200, AtMg00650, and AtMg00270, respectively), we could identify all Complex I subunits in our proteomic data set. Additionally, we identified eight previously proposed assembly factors (Meyer et al., 2019), five γ-carbonic anhydrases, and five additional proteins proposed to form a matrix-exposed domain attached to Complex I (Sunderhaus et al., 2006). Complex II is composed of eight subunits (Millar et al., 2004). In addition to its function in the OXPHOS pathway, Complex II also participates in the tricarboxylic acid cycle (TCA). Six out of eight subunits of Complex II and the assembly factor SDHAF2 were identified in our data set. Additionally, peptides of all proteins and isoforms of Complex III, eight of its assembly factors, and proteins previously defined as alternative pathways (Meyer et al., 2019) were identified in our data set. Complex V consists of 15 subunits, of which 13 were identified in our proteomic data set; Atp6 (AtMg00410 and AtMg011701) and Atp9 (AtMg01080) were the only two subunits that could not be identified. In addition, we found three of the five proposed assembly factors (Meyer et al., 2019). The cytochrome c oxidase complex (Complex IV) consists of 16 proposed subunits in Arabidopsis (Mansilla et al., 2018). Of these, 9 subunits and 11 assembly factors of Complex IV were identified in our proteomic data set. Furthermore, we identified three proteins involved in the assembly of OXPHOS supercomplexes (Meyer et al., 2019).

Except the abovementioned subunits of the SDH complex, all proteins of the TCA cycle and GDC multienzyme system were identified in the rapidly isolated mitochondria. Additionally, our proteomic data set contained a number of pentatricopeptide and tetratricopeptide repeat proteins involved in RNA metabolism as well as heat shock proteins and ribosomes involved in protein control and turnover (Supplemental Table S1, Supplemental Table S2).

In Arabidopsis, the majority of mitochondrial proteins are encoded by nuclear genes, translated in the cytosol, and then imported into the mitochondria. The import and sorting of nuclear-encoded mitochondrial preproteins requires functional TOM and sorting and assembly machinery (SAM) in the OMM, mitochondrial IMS import and assembly (MIA) machinery in the IMS, and translocase of the IMM (TIM) in the IMM (Murcha et al., 2014). All mitochondrial preproteins enter the mitochondria via the TOM complex in the OMM. In the IMS, membrane proteins are sorted via the small TIM proteins toward the SAM or TIM22 complex for incorporation into the OMM or IMM. Soluble proteins of the mitochondrial matrix are imported via the TIM17:23 complex, and proteins that remain in the IMS are processed via the MIA machinery (Murcha et al., 2014). In this study, we identified all of these proteins in our proteome data set, except the OMM protein TOM6, IMS protein ERV1, IMM protein PRAT5, and matrix proteins MGE1 and ZIM17. Additionally, we identified plant-specific import components, including OM64, PRAT3, and PRAT4, in the OMM (Murcha et al., 2015) and proteins of the mitochondrial contact site and cristae organizing system (MICOS), which connects the IMM to OMM (van der Laan et al., 2016). Other notable OMM proteins identified in our proteomic data set included GTPases MIRO1 and MIRO2, lipid biosynthesis protein PECT, and β-barrel proteins VDAC1–4 (Supplemental Table S1, Supplemental Table S2). Recently, it was shown that the cytosolic protein GAPC interacts with VDAC (Schneider et al., 2018); we also identified GAPC in our proteomic data set.

Overall, we conclude that mitochondria isolated using our rapid isolation method are suitable for proteomic analyses.

### Differential analysis of the mitochondrial proteome of transgenic Col-0 and *bou-2* lines

To assess the effect of *bou-2* mutation on the mitochondrial proteome, we performed comparative proteomic analysis of transgenic Col-0 and *bou-2* lines. A total of 47 proteins showed significantly increased abundance in the mutant, of which five were localized to the mitochondria. Additionally, 44 proteins showed significantly decreased abundance in the transgenic *bou-2* samples, of which 22 were predicted to be localized to the mitochondria (Table 1); among these proteins, BOU was the least abundant. The *bou-2* line is a GABI-Kat line that carries a T-DNA insertion in the second exon of the *BOU* gene (Kleinboelting et al., 2012; Eisenhut et al., 2013). We identified two peptides of BOU in at least three of the four replicates of transgenic *bou-2* samples. Both peptides were translated from the first exon of the gene. Because the T-DNA was inserted in the second exon of the gene, it is possible that the first exon was translated. However, a functional protein is not synthesized in the knockout mutant (Eisenhut et al., 2013). Among the mitochondrial proteins showing significantly reduced abundance in transgenic *bou-2* seedlings, we identified six proteins of the OXPHOS pathway (three Complex I proteins and one protein each of Complex II, III, and V), two proteins involved in protein translocation, two proteins involved in metabolite transport, two proteins involved in lipid metabolism, three proteins involved in protein turnover/synthesis, one protein involved in the TCA cycle, and six proteins (including FDH) involved in other processes. Among the proteins with significantly increased abundance in transgenic *bou-2* mutant, we identified two proteins involved in RNA/DNA metabolism, one protein involved in THF metabolism, one MIRO-related GTPase, and one LETM1-like protein (Table 1).

**Table 1:** List of significant differences in protein abundance between Col-0 *UB10p-3×HA-sGFP-TOM5* and *bou-2 UB10p-3×HA-sGFP-TOM5*. Difference was calculated as change of log_2_ of normalized intensity. List includes only proteins that show mitochondrial localization. List ranges from most downregulated in *bou-2 UB10p-3×HA-sGFP-TOM5* (top) to most upregulated *bou-2 UB10p-3×HA-sGFP-TOM5* (bottom). Significance was calculated with Student’s t-test, *P* < 0.05.

| AGI | Gene symbol | Gene description | Difference |
| --- | --- | --- | --- |
| AT5G46800 | BOU | Mitochondrial substrate carrier family protein | -8,66407 |
| AT2G42310 | AT2G42310 | ESSS subunit of NADH:ubiquinone oxidoreductase (complex I) protein | -2,0953 |
| AT5G41685 | AT5G41685 | Mitochondrial outer membrane translocase complex, subunit Tom7 | -1,90711 |
| AT3G03100 | AT3G03100 | NADH:ubiquinone oxidoreductase, 17.2kDa subunit | -1,33356 |
| AT5G53650 | AT5G53650 | ABC transporter A family protein | -1,25252 |
| AT5G67590 | FRO1 | NADH-ubiquinone oxidoreductase-like protein | -1,0968 |
| AT5G40810 | AT5G40810 | Cytochrome C1 family | -1,06444 |
| AT3G27280 | PHB4 | prohibitin 4 | -0,94888 |
| AT3G27380 | SDH2-1 | succinate dehydrogenase 2-1 | -0,764682 |
| AT4G37660 | AT4G37660 | Ribosomal protein L12/ ATP-dependent Clp protease adaptor protein ClpS family protein | -0,732468 |
| AT2G42210 | OEP16-3 | Mitochondrial import inner membrane translocase subunit Tim17/Tim22/Tim23 family protein | -0,710023 |
| AT3G55400 | OVA1 | methionyl-tRNA synthetase / methionine-tRNA ligase / MetRS (cpMetRS) | -0,705525 |
| AT5G14780 | FDH | formate dehydrogenase | -0,697987 |
| AT2G38670 | PECT1 | phosphorylethanolamine cytidyltransferase 1 | -0,606253 |
| AT5G63400 | ADK1 | adenylate kinase 1 | -0,603703 |
| AT1G79230 | MST1 | mercaptopyruvate sulfurtransferase 1 | -0,587758 |
| AT4G30010 | AT4G30010 | ATP-dependent RNA helicase | -0,545131 |
| AT3G03420 | AT3G03420 | Ku70-binding family protein | -0,531796 |
| AT4G31810 | AT4G31810 | ATP-dependent caseinolytic (Clp) protease/crotonase family protein | -0,461768 |
| AT1G19140 | AT1G19140 | ubiquinone biosynthesis COQ9-like protein | -0,455841 |
| AT4G31460 | AT4G31460 | Ribosomal L28 family | -0,430804 |
| AT1G54220 | AT1G54220 | Dihydrolipoamide acetyltransferase, long form protein | -0,324755 |
| AT3G59820 | LETM1 | LETM1-like protein | 0,42657 |
| AT5G27540 | MIRO1 | MIRO-related GTP-ase 1 | 0,447933 |
| AT3G10160 | DFC | DHFS-FPGS homolog C | 0,507662 |
| AT1G71260 | ATWHY2 | WHIRLY 2 | 0,51931 |
| AT5G15980 | AT5G15980 | Pentatricopeptide repeat (PPR) superfamily protein | 0,8335 |

Previously, Eisenhut and colleagues showed that GDC activity is reduced in mitochondria isolated from 4-week-old *bou-2* mutant plants (Eisenhut et al., 2013). The authors showed that the *bou-2* mutant accumulated higher amounts of glycine than the wild type and exhibited differential amount and status of the P-protein. Immunoblot analysis showed no differences in the levels of other GDC proteins in the *bou-2* mutant compared with the wild type (Eisenhut et al., 2013). In our proteomic data set, none of the proteins of the GDC complex or SHMT showed significant differences between transgenic Col-0 and *bou-2* seedlings (Table 2). However, the amounts of GLDP1, GLDP2, GDCH1, GDCL1, GDCL2, and GLDT were slightly reduced, whereas those of SHM1, SHM2, and GDCH3 were increased in the mutant compared with Col-0. Among these proteins, the strongest reduction was detected in the amount of GDCH1. However, differences in protein levels between transgenic Col-0 and *bou-2* seedlings were not statistically significant. Only one peptide related to GDCH2 was detected in our data set; however, because it was detected in only one of the four replicates, it is not listed Supplemental Table S1.

**Table 2:**
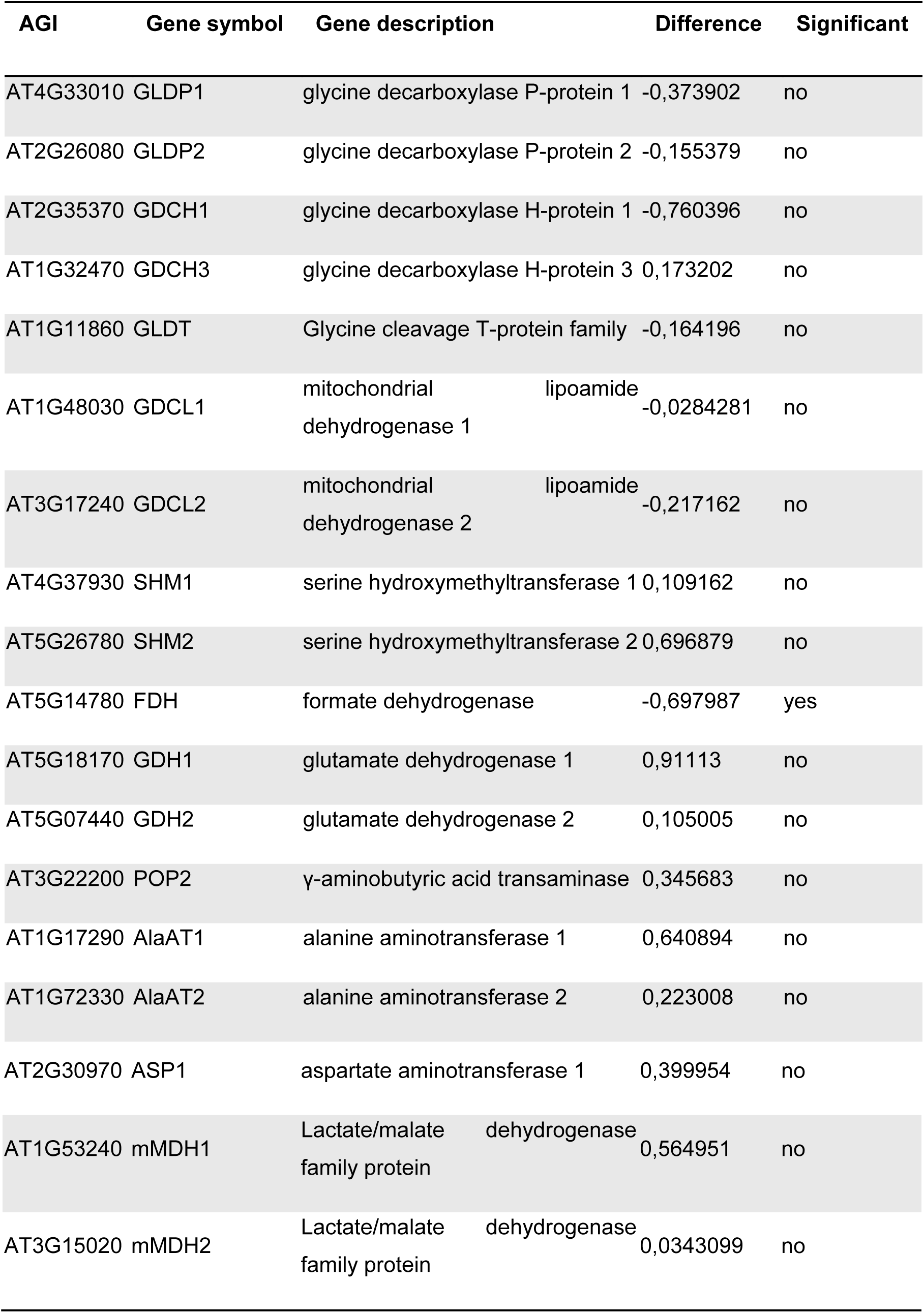
List of changes in protein abundance of glycine decarboxylase proteins, serine hydroxymethyltransferase (SHM) and the enzymes malate dehydrogenase (MDH), formate dehydrogenase (FDH), aspartate aminotransferase (ASP), γ-aminobutyric acid transaminase (POP2), alanine aminotransferase (AlaAT) and glutamate dehydrogenase (GDH). Difference was calculated as change of log_2_ of normalized intensity. Significance was calculated with Student’s t-test, *P* < 0.05.

In this study, we showed that mitochondria of the *bou-2* mutant displayed reduced AlaAT activity, increased GluDH activity, and no change in MDH, AspAT, GABA-T, and FDH activities under elevated CO_2_ conditions compared with that under ambient CO_2_ conditions (Fig. 4A–F). Except FDH, none of the assayed enzymes showed significantly altered amounts in our proteomic data set (Table 2). The amount of FDH was significantly reduced in transgenic *bou-2* samples; however, its activity was not altered in the mutant under both elevated and ambient CO_2_ conditions, indicating post-translational modification of FDH. The level of AlaAT was slightly increased in the mutant but showed only 60% activity compared with Col-0 under both elevated and ambient CO_2_ conditions. The activities of MDH, AspAT, and GABA-T did not differ between transgenic Col-0 and *bou-2* mutant under elevated CO_2_ conditions, although these proteins were more abundant in transgenic mutant samples. The activity of GluDH was significantly increased in the mutant compared with Col-0 under elevated CO_2_ conditions, which may be associated with the increased amount of protein detected in the transgenic *bou-2* seedlings in our proteomic data set. However, this increase was not statistically significant.

Overall, we conclude that differences in the activities of MDH, AspAT, GluDH, AlaAT, GABA-T, and AlaAT measured in this study and that of GDC measured in a previous study most likely do not relate to changes in protein abundance in mitochondria of Col-0 vs. *bou-2* mutant but instead might be caused by metabolic impairment or post-translational modifications.

## DISCUSSION

Recently, analyses of mitochondrial proteome content, complexome composition, post-translational modifications, energy metabolism, OXPHOS complex formation and function, protein translocation, and metabolite shuttles have been conducted to further our understanding of mitochondrial metabolism in Arabidopsis (König et al., 2014; Fromm et al., 2016; De Col et al., 2017; Rao et al., 2017; Senkler et al., 2017; Porcelli et al., 2018; Hu et al., 2019; Kolli et al., 2019; Meyer et al., 2019; Nickel et al., 2019). Many of these analyses required the isolation of intact mitochondria. Here, we report a procedure for the rapid isolation of HA-tagged mitochondria from transgenic Arabidopsis lines via co-IP. Mitochondria isolated using this method showed high enrichment of mitochondrial marker proteins, with only minor contamination, as demonstrated by immunoblot and quantitative proteomic analyses (Fig. 3, Supplemental Table S1, Supplemental Table S2).

The method reported here enables the isolation of intact mitochondria from Arabidopsis seedlings in less than 25 min (Fig. 2, Fig. 3). Moreover, by omitting the first centrifugation step, the isolation time could be shortened to 18 min, although the resulting mitochondrial fraction contained a higher level of other contaminating cellular components. The mitochondrial fraction used for proteomic analyses in this study was obtained using the slightly longer protocol that results in lower contamination with non-mitochondrial proteins. Nevertheless, this isolation method is significantly faster than the standard isolation procedures that generally take up to several hours. To date, we have been able to successfully isolate mitochondria from whole seedlings, leaves, and roots (data not shown). Moreover, this method could be used for the rapid isolation of mitochondria from other plant tissues such as flowers and developing seeds because (1) the expression of the affinity tag is driven by the ubiquitous *UB10p* promoter (Supplemental Fig. S1) and (2) only a small amount of starting material (as low as 1 g) is needed. Additionally, this method is advantageous for the isolation of mitochondria from very young tissues or mutants with growth defects or reduced biomass accumulation. The rapid isolation method is superior to the standard isolation protocols with respect to the yield of mitochondria; we were able to isolate 200 µg of total mitochondrial protein from 1 g of starting material and up to 700 µg total mitochondrial protein from 10 g of whole Arabidopsis seedlings. By contrast, the standard isolation protocols yield only 1.2 mg mitochondria from 50 g of leaves (Keech et al., 2005). Generally, a higher yield is expected using our isolation method, as the HA-tag has high affinity for its cognate antibody. Further optimization of the protocol and bead to extract ratio may result in even higher yields.

The purity of affinity-purified mitochondria was similar to that of mitochondria isolated using standard protocols including density gradients (Senkler et al., 2017). Immunoblot analysis revealed only minor contamination with proteins of the ER or peroxisomes in affinity-purified mitochondria (Fig. 3). These results were corroborated by quantitative proteomic analysis. Less than 3% of all quantified proteins were assigned to peroxisomes, ER, Golgi, vacuoles, endomembranes, plasma membranes, nuclei, and cytosol. The highest contamination was due to plastid-localized proteins (Supplemental Table S1, Supplemental Table S2). However, we cannot exclude the possibility that some of the contaminants bound non-specifically to the beads, despite extensive washing. Approximately 80% of the identified proteins and 90% of the identified peptides showed mitochondrial localization (Supplemental Table S1, Supplemental Table S2). Thus, we conclude that the purity of mitochondria isolated using the rapid affinity purification method is comparable with that of mitochondria isolated using traditional methods. We cannot comment on the physiological activity and coupling state of affinity-purified mitochondria because we could not efficiently elute the mitochondria from the magnetic beads with HA peptide. However, mitochondria could be efficiently eluted using SDS-PAGE or detergent lysis buffer. Elution of intact mitochondria might be achievable by integrating a protease cleavage site between the HA-tag and GFP; this will be explored in future studies.

In our data set, we were able to identify 96% and 99% of the proteins detected by Klodmann et al. (2011) and Senkler et al. (2017), respectively, in previous analyses of mitochondrial proteomes and their complexome composition (Supplemental Table S1, Supplemental Table S2). On the basis of a recent review on the composition and function of OXPHOS in mitochondria (Meyer et al., 2019), we were able to identify 86% of all the predicted subunits and assembly factors. To date, all of the proteins reviewed in Meyer et al. (2019) have not been confirmed, and it might require an in-depth analysis or membrane enrichment to confirm their presence in mitochondria in proteomic studies. Previously, it was reported that the subunit ND4L is difficult to detect in proteomic studies because of its hydrophobic nature (Peters et al., 2013). Consistent with this observation, we also could not identify this protein in our proteome or in the list of quantified peptides (Supplemental Table S1, Supplemental Table S2). However, we were able to detect all proteins of the TCA cycle (except two subunits of Complex II), all GDC proteins, majority of the proteins and subunits of the TIM/TOM protein import apparatus, metabolite transporters of amino acids, dicarboxylic acids, cofactors, ions and energy equivalents (e.g., BOU, BAC, UCP, DCT, SAMC, NDT, APC, AAC, and PHT), as well as many proteins involved in protein synthesis/turnover and DNA/RNA metabolism (Supplemental Table S1, Supplemental Table S2). Thus, our rapidly isolated mitochondria showed good purity, integrity, and functionality.

Among all of the identified proteins, 20% could not be assigned to mitochondria. These proteins included components of PSI and PSII; these could be clearly categorized as contamination. However, we also identified proteins with unknown function and no clear prediction of localization. In addition, we detected proteins such as GAPC, which was previously shown to interact with the OMM protein VDAC (Schneider et al., 2018). Therefore, we predict that some of the proteins classified as contaminants might represent novel mitochondrial proteins, for example, as part of OMM protein complexes in the cytosol or as components of complexes at organellar contact sites. However, this needs to be evaluated in more detail in the future.

Application of the novel method to a mutant lacking the mitochondrial glutamate transporter BOU resulted in the detection of surprisingly few changes in protein abundances compared with Col-0. Only 22 mitochondrial proteins showed a significant reduction in abundance in the *bou-2* mutant, whereas five proteins were significantly increased in abundance. However, only changes in a mitochondrial folylpolyglutamate synthetase (FGPS) and in FDH, which contributes to the production of CO_2_ by oxidizing formate, might be connected to photorespiration. Formate is released from 10-formyl-THF by 10-formyl deformylase, an enzyme involved in the maintenance of the THF pool in mitochondrial matrix (Collakova et al., 2008). Reduced GDC activity in *bou-2* might result in a lower production of formate and finally a reduced abundance of FDH. FGPS is involved in vitamin B9 metabolism by catalyzing the glutamylation of THF, a cofactor of GLDT and SHM (Hanson and Gregory, 2011). No significant changes were detected in the abundance of any of the proteins of the GDC multienzyme system. SHM1, SHM2, and GDCH3 were slightly more abundant in the mutant than Col-0, whereas the others seem to be slightly reduced; however, this trend did not meet the significance threshold (Table 2). The observed changes were not pronounced and did not explain why the GDC activity was reduced to approximately 15% of the Col-0 level in the *bou-2* mutant. A possible explanation could be that Eisenhut and coworkers used 4-week-old rosettes (Eisenhut et al., 2013), whereas we used 10-day-old seedlings in the current study. This possibility is supported by the observation that no difference in MDH activity could be detected between 10-day-old mutant and Col-0 seedlings (Fig. 5A). However, in 4-week-old leaves, the MDH activity was significantly reduced in the mutant compared with Col-0 (Supplemental Fig. S2). In contrast to GDC, the abundance of FDH was significantly reduced in the mutant, whereas its activity was unaltered compared with Col-0 (Table 1, Fig. 5B). Previously, FDH was identified in a lysine acetylome study of Arabidopsis mitochondria from 10-day-old Col-0 seedlings (König et al., 2014). This might indicate that FDH activity is more likely regulated by post-translational modifications than by protein abundance.

The BOU protein was recently assigned the function of a glutamate transporter (Porcelli et al., 2018). Glutamate is indirectly linked to photorespiration, as it is needed for the polyglutamylation of THF, which increases its stability and promotes the activity of THF-dependent enzymes (Hanson and Gregory, 2011). However, in addition to BOU, mitochondria-localized uncoupling proteins 1 and 2 show glutamate uptake activity in Arabidopsis (Monné et al., 2018). This raises the question why knockout of the *BOU* gene leads to a photorespiratory phenotype in young tissues, if BOU is not the only glutamate transporter of mitochondria. Additionally, polyglutamylation is not restricted to mitochondria, as Arabidopsis contains three isoforms of FGPS localized to the mitochondria, plastids, and cytosol (Hanson and Gregory, 2011). Folate transporters prefer monoglutamylated forms of THF, whereas enzymes generally prefer polyglutamylated forms (Suh et al., 2001). These data indicate that BOU performs other functions, in addition to glutamate transport *in vivo*. We exclude its function as a mitochondrial glutamate/glutamine shuttle, as BOU shows no glutamine uptake activity (Porcelli et al., 2018). However, it is possible that BOU is involved in folate metabolism, as folate biosynthesis occurs only in mitochondria (Hanson and Gregory, 2011). This possibility, however, needs to be investigated in future studies.

## CONCLUSIONS

Our experiments show that affinity-tagging is a powerful tool not only for the analysis of protein-protein interactions but also for the isolation of functional organelles from Arabidopsis. The mitochondria isolated using this method showed high purity and integrity. Future studies will be required to assess the physiological state of the isolated mitochondria after elution from magnetic beads and to determine the applicability of this method for metabolite analyses. To conduct metabolite analyses, it is encouraging that the LC/MS-compatible buffer system developed previously for mammalian mitochondria (Chen et al., 2016) can also be used for the isolation of mitochondria from plant tissues. Expressing the affinity tag under the control of cell-specific promoters will allow the isolation of mitochondria from specific cell types, such as meristems or guard cells (Yang et al., 2008; Schürholz et al., 2018). The use of cell-specific promoters in our construct will help unravel the complex role of mitochondria in various cell types. We expect that similar tagging strategies will be applicable to other plant cell organelles, such as plastids and peroxisomes. Moreover, simultaneous expression of several different affinity tags will facilitate the affinity purification of different organelles from a single extract.

## MATERIAL AND METHODS

### Pant growth conditions

Arabidopsis ecotype Col-0 and *bou-2* mutant (GABI-Kat line number 079D12; http://www.gabi-kat.de/db/lineid.php) (Kleinboelting et al., 2012) were used in this study. Seeds were sterilized by washing with 70% (v/v) ethanol supplemented with 1% (v/v) Triton X-100 twice for 10 min each, followed by washing with 100% ethanol twice for 10 min each. Seeds were grown on half-strength Murashige and Skoog (1/2 MS) medium (pH 5.8) supplemented with 0.8% (w/v) agar. Seeds were subjected to cold stratification for 2 days at 4°C. After germination, seedlings were grown under 12 h light/12 h dark photoperiod under 100 µmol m^−2^ s^−1^ light intensity and 3,000 ppm CO_2_-enriched atmosphere, unless otherwise stated.

### Construction of transgenic lines

The HA epitope-tag was amplified with a start codon from the Gateway binary vector pGWB15. The coding sequence (CDS) of sGFP, minus the start and stop codons but including a linker peptide (GGSG) at the 5’ and 3’ ends, was amplified from the Gateway binary vector pGWB4. The CDS of *TOM5* (minus the start codon) was amplified from Arabidopsis cDNA. Starting from the 5’-end to the 3’-end, the amplified 3×HA-tag, sGFP, and TOM5 were cloned into the pUTKan vector under the control of the Arabidopsis *UB10p* using restriction endonucleases. The construct was introduced into *Agrobacterium tumefaciens*, strain GV3101::pMP90, which was then introduced into Col-0 and *bou-2* plants via *Agrobacterium*-mediated transformation using the floral dip method, as described previously (Clough and Bent, 1998).

### Confocal laser scanning microscopy

The expression of 3×HA-sGFP-TOM5 was verified via confocal laser scanning microscopy using the Zeiss LSM 78 Confocal Microscope and Zeiss ZEN software. The Col-0 and *bou-2* seedlings regenerated from independent transformation events were incubated with 200 nM MitoTracker Red CMXRos (Molecular Probes) in 1/2 MS supplemented with 3% (w/v) sucrose for 15 min. Images were captured using the following excitation/emission wavelengths: sGFP (488 nm/490–550 nm) and MitoTracker Red CMXRos (561 nm/580–625 nm). Pictures were processed using the ImageJ software (https://imagej.nih.gov/ij/).

### Rapid isolation of mitochondria using co-IP

Epitope-tagged mitochondria were rapidly isolated using co-IP, as described previously (Chen et al., 2016). Briefly, 1–10 g of Arabidopsis seedlings were harvested and homogenized in KPBS (10 mM KH_2_PO_4_ [pH 7.25] and 136 mM KCl) using a Warren blender. The resulting homogenate was filtered through three layers of miracloth supported by a nylon mesh and centrifuged at 2,500 × *g* for 5 min. The pellet containing cell debris and chloroplasts was discarded. The supernatant was subsequently centrifuged at 20,000 × *g* for 9 min. The pellet representing the crude mitochondrial fraction was resuspended in KPBS using a fine paintbrush and homogenized using a Potter-Elvehjem. Crude mitochondria were incubated with pre-washed magnetic anti-HA beads (ThermoFisher Scientific) on an end-over-end rotator for 5 min. Magnetic beads were separated using a magnetic stand and washed at least three times with KPBS. In the detergent treatment control, KPBS was supplemented with 1% (v/v) Triton X-114 in all washing steps. The purified mitochondria were lysed using mitochondria lysis buffer (50 mM TES/KOH [pH 7.5], 2 mM EDTA, 5 mM MgCl_2_, 10% [v/v] glycerol, and 0.1% [v/v] Triton X-100) for enzyme activity assays and immunoblot analysis or directly frozen in liquid nitrogen for proteome and metabolite analyses. All steps were carried out at 4°C. The amount of protein recovered after lysis was determined using the Quick Start™ Bradford Protein Assay Kit (Bio-Rad), with bovine serum albumin (BSA) as the standard.

### Isolation of mitochondria using the traditional approach

Mitochondria were isolated from 10-day-old Col-0 seedlings via differential centrifugation and Percoll gradient purification, as described previously (Kühn et al., 2015).

### Immunoblot analysis

25 µg of total leaf extract and 6.45 µg of isolated mitochondria fractions were heated at 96°C in SDS-PAGE loading buffer for 10 min and separated on 12% SDS-polyacrylamide gels (Laemmli, 1970). Proteins were transferred to 0.2 µm polyvinylidene difluoride membranes (PVDF) or 0.45 µm nitrocellulose membranes using standard protocols. Protein transfer was verified by staining the membranes with Ponceau S red. Membranes were blocked according to the manufacturer’s instructions for 1 h, washed with Tris-buffered saline containing 0.1% (v/v) Tween-20 (TBST) and subsequently incubated with either a primary antibody or a single-step antibody overnight at 4°C. Antibodies against marker proteins were diluted as follows: anti-AOX (1:1,000), anti-IDH (1:5,000), anti-VDAC1 (1:5,000), anti-HA-HRP (1:5,000), anti-RbcL (1:7,500), anti-Cat (1:1,000), anti-BiP2 (1:2,000), anti-Histone H3 (1:5,000), and anti-HSC70 (1:3,000). Membranes were washed with TBST twice for 10 min each and incubated with the secondary goat anti-Rabbit-HRP antibody (1:5,000) at room temperature for 1 h or at 4°C overnight. Subsequently, membranes were washed with TBST five times for 5 min each and visualized using a chemiluminescence detection system (Immobilon Western HRP Substrate, Merck Millipore). All steps were carried out with phosphate-buffered saline (PBS), when using anti-Cat antibody.

### Enzyme assays

Activities of mitochondrial enzymes were measured using a spectrophotometer, based on the absorbance at 340 nm. The activity of MDH was measured based on the oxidation of NADH to NAD^+^ at 340 nm, as described previously (Tomaz et al., 2010). The reaction mixture contained 50 mM KH_2_PO_4_ (pH 7.5), 0.2 mM NADH, 5 mM EDTA, 10 mM MgCl_2_, 2 mM OAA (Tomaz et al., 2010), and 0.1–0.4 µg total mitochondrial protein. AspAT activity was measured in a reaction coupled with MDH, as described previously (Wilkie and Warren, 1998), with 0.3–1 µg total mitochondrial protein per assay. No external pyridoxal-5’-phosphate was added to the reaction mixture. GluDH activity was measured as described previously (Turano et al., 1996), based on the reduction of NAD^+^ to NADH at 340 nm. To determine the amination activity, 0.5–2 µg total mitochondrial protein was used per assay. GABA-T activity was measured in a reaction coupled with succinate-semialdehyde dehydrogenase (SSADH), as described previously (Clark et al., 2009). The assay buffer contained 50 mM TAPS (pH 9), 0.2 mM NAD^+^, 0.625 mM EDTA, 8 mM GABA, 2 mM pyruvate, 1 U/ml SSADH, and 2–5 µg total mitochondrial protein. The NAD^+^-dependent SSADH was purified from *Escherichia coli*, as described previously (Clark et al., 2009). The recombinant purified protein catalyzed the production of NADH with a specific activity of 1.6 U/mg protein. AlaAT activity was measured in reaction coupled with lactate dehydrogenase, as described previously (Miyashita et al., 2007), with 1–5 µg total mitochondrial protein. FDH activity was measured based on the reduction of NAD^+^ to NADH at 340 nm and 30°C, with 1–5 µg total mitochondrial protein. The assay buffer contained 100 mM KH_2_PO_4_ (pH 7.5), 1mM NAD^+^, and 50 mM sodium formate.

### Sample preparation for LC/MS analysis

To elute proteins from magnetic beads, 30 µl of Laemmli buffer was added to the reaction mixture, and samples were incubated at 95°C for 10 min. Subsequently, 20 µl of protein sample was loaded on an SDS-polyacrylamide gel for in-gel digestion. The isolated gel pieces were reduced using 50 µl of 10 mM DTT, then alkylated using 50 µl of 50 mM iodoacetamide, and finally digested using 6 µl of trypsin (200 ng) in 100 mM ammonium bicarbonate. The peptides were resolved in 15 µl of 0.1% trifluoracetic acid and subjected to LC/MS analysis.

### LC/MS analysis

The LC/MS analysis was performed on a Q Exactive Plus mass spectrometer (Thermo Scientific, Bremen, Germany) connected to an Ultimate 3000 Rapid Separation LC system (Dionex; Thermo Scientific, Idstein, Germany) and equipped with an Acclaim PepMap 100 C18 column (75 µm inner diameter × 25 cm length × 2 mm particle size; Thermo Scientific, Bremen, Germany). The length of the isocratic LC gradient was 120 min. The mass spectrometer was operated in positive mode and coupled with a nano electrospray ionization source. Capillary temperature was set at 250°C, and source voltage was set at 1.4 kV. The survey scans were conducted at a mass to charge (m/z) ranging from 200 to 2000 and a resolution of 70,000. The automatic gain control was set at 3,000,000, and the maximum fill time was set at 50 ms. The ten most intensive peptide ions were isolated and fragmented by high-energy collision dissociation (HCD).

### Computational MS data analysis

Peptide and protein identification and quantification was performed using MaxQuant version 1.5.5.1 (MPI for Biochemistry, Planegg, Germany) with default parameters. The identified Arabidopsis peptides and proteins were queried against a specific proteome database (UP0000006548, downloaded 12/11/17) from UniProt. The oxidation and acetylation of methionine residues at the N-termini of proteins were set as variable modifications, while carbamidomethylations at cysteine residues were considered as fixed modification. Peptides and proteins were accepted with a false discovery rate of 1%. Unique and razor peptides were used for label-free quantification, and peptides with variable modifications were included in the quantification. The minimal ratio count was set to two, and the ‘matched between runs’ option was enabled.

Normalized intensities, as provided by MaxQuant, were analyzed using the Perseus framework (version 1.5.0.15; MPI for Biochemistry, Planegg, Germany). Only proteins containing at least two unique peptides and a minimum of three valid values in at least one group were quantified. Proteins which were identified only by site or marked as a contaminant (from the MaxQuant contaminant list) were excluded from the analysis. Differential enrichment of proteins in the two groups (*Col-0*; *bou-2*) was assessed using Student’s *t*-test. Significance analysis was applied on log_2_-transformed values using an S0 constant of 0 and a false discovery rate of 5%, as threshold cutoffs.

The MS proteomics data has been deposited to the ProteomeXchange Consortium via the PRIDE partner repository with the data set identifier PXD014137.

## SUPPLEMENTAL MATERIAL

**Supplemental Table S1:** List of proteins identified and quantified in Col-0 and *bou-2* plants expressing the 3×HA-sGFP-TOM5 protein.

**Supplemental Table S2:** List of raw intensities and reliability of all quantified peptides.

**Supplemental Figure S1:** Schematic representation of the vector used to label mitochondria with triple HA-tag.

**Supplemental Figure S2:** Activity of malate dehydrogenase (MDH) in mitochondria rapidly isolated from 4-week-old Col-0 and *bou-2* leaves expressing the *UB10p-3×HA-sGFP-TOM5* construct.

## ACKNOWLEDGEMENTS

This work was supported by the Deutsche Forschungsgemeinschaft (CRC 1208 and funding under Germany’s Excellence Strategy – EXC-2048/1 – project ID 390686111”).

## SUPPLEMENTARY DATA

**Supplemental Table S1: List of proteins identified and quantified in Col-0 and *bou-2* plants expressing the 3×HA-sGFP-TOM5 protein.** Intensities of identified proteins are given as log_2_-transformed values. Subcellular localization was assigned using SUBA4. Difference between Col-0 and *bou-2* was calculated as the change in log_2_-transformed values. Significance was calculated using the Student *t*-test (*P* < 0.05).

**Supplemental Table S2: List of raw intensities and reliability of all quantified peptides.** PEP: Posterior Error Probability of identification.

**Supplemental Figure S1: Schematic representation of the vector used to label mitochondria with triple HA-tag.** The N-terminal end of Arabidopsis gene encoding the outer mitochondrial membrane protein TOM5 (At5g08040) was fused to the *synthetic green fluorescent protein* (*sGFP*) gene labeled with a triple hemagglutinin (3×HA) tag for co-Immunopurifiaction. The cassette was cloned into the pUTKan vector under the control of the Arabidopsis *UBIQUITIN10* promoter (*UB10p*) and stably introduced into Arabidopsis via the floral dip method.

**Supplemental Figure S2: Activity of malate dehydrogenase (MDH) in mitochondria rapidly isolated from 4-week-old Col-0 and *bou-2* leaves expressing the *UB10p-3×HA-sGFP-TOM5* construct.** MDH activity was measured as described in Material and Methods. Activities were calculated from initial slopes. Enzyme activity in transgenic Col-0 was set to 100%. Data represent mean ± SD of three biological replicates. Asterisks indicate statistically significant differences (***, *P* < 0.001; Student’s *t*-test).

