## Supplementary figures and images for "Rapid single-step affinity purification of HA-tagged mitochondria from *Arabidopsis thaliana*"

### Supplemental Figures 1-2

**Supplemental Figure S1**

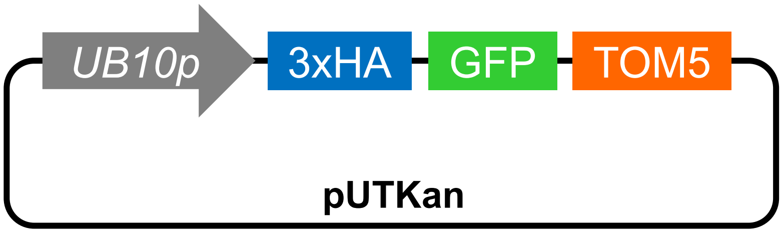

**Supplemental Figure S2**

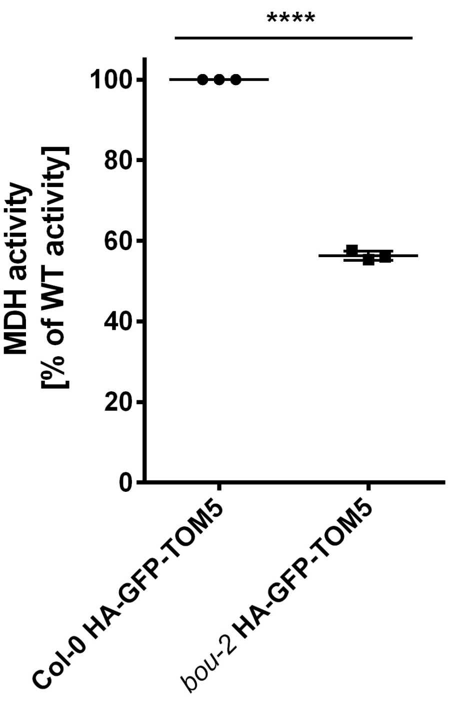
